# Turning the Stimulus On and Off Dynamically Changes the Direction of Alpha Traveling Waves

**DOI:** 10.1101/2020.04.15.041756

**Authors:** Zhaoyang Pang, Andrea Alamia, Rufin VanRullen

## Abstract

Traveling waves have been studied to characterize the complex spatiotemporal dynamics of the brain. Several studies have suggested that the propagation direction of alpha traveling waves can be task-dependent. For example, a recent EEG study from our group found that forward waves (i.e. occipital to frontal, FW waves) were observed during visual processing, whereas backward waves (i.e. frontal to occipital, BW waves) mostly occurred in the absence of sensory input. These EEG recordings, however, were obtained from different experimental sessions and different groups of subjects. To further examine how the waves’ direction changes between task conditions, 13 participants were tested on a target detection task while EEG signals were recorded simultaneously. We alternated visual stimulation (5 s display of visual luminance sequences) and resting state (5 s of black screen) within each single trial, allowing us to monitor the moment-to-moment progression of traveling waves. As expected, the direction of alpha waves was closely linked with task conditions. First, FW waves from occipital to frontal regions, absent during rest, emerged as a result of visual processing, while BW waves in the opposite direction dominated in the absence of visual inputs, and were reduced (but not eliminated) by external visual inputs. Second, during visual stimulation (but not rest), both waves coexisted on average, but were negatively correlated. In summary, we conclude that the functional role of alpha traveling waves is closely related with their propagating direction, with stimulus-evoked FW waves supporting visual processing and spontaneous BW waves involved more in top-down control.

## Introduction

Neural oscillations at various temporal frequencies are ubiquitous in the human brain, and in the spatial domain, an increasing number of studies suggest that these oscillations could be organized as traveling waves across brain regions. The existence of traveling waves has been reported across multiple species (Ermentrout & Kleinfeld, 2001; Sato et al., 2012), at differing scales of measurements (Muller et al., 2018), and under various stimulation conditions (Nauhaus et al., 2012; Sato et al., 2012). Since the propagation of the traveling waves covers highly distributed brain regions, researchers have attempted to relate their functional significance to various aspects of the traveling waves. In particular, the directionality of traveling waves is believed to be functionally relevant (Bahramisharif et al., 2013; Fellinger et al., 2012; Klimesch, Hanslmayr, et al., 2007; Patten et al., 2012). For example, Halgren and colleagues showed that, during wakefulness with open or closed eyes, alpha oscillations recorded with intra-cortical electrodes from epilepsy patients propagated from antero-superior cortex towards postero-inferior occipital poles (Halgren et al., 2019). However, in another intra-cortical study (Zhang et al., 2018), when subjects were instructed to complete a visual memory task, traveling waves in the theta-alpha band (2-15 Hz) propagated from posterior to anterior brain areas. This apparent forward direction of traveling waves was also reported in studies of so-called “perceptual echoes”, which constitute a direct index of sensory processing (VanRullen & Macdonald, 2012). Participants were stimulated with random (white-noise) luminance sequences, and the resulting impulse response function showed a long-lasting 10Hz oscillation (or perceptual echo); importantly, the spatial distribution of echo phase was organized as a traveling wave propagating from posterior to frontal sensors (Lozano-Soldevilla & VanRullen, 2019; Alamia & VanRullen, 2019). It thus seems that the directionality of traveling waves could be task-dependent. To clarify the traveling direction with respect to various experimental conditions, a recent study (Alamia & VanRullen, 2019) from our group simulated alpha oscillations as a cortical traveling wave within a predictive coding framework. The predictive coding framework characterizes a hierarchical network where higher levels of brain regions predict the activity of lower levels, and the unexplained residuals (i.e., prediction errors) are passed back to higher layers. The study revealed that the recursive nature of predictive coding not only gave rise to alpha oscillations but also explained their propagating dynamics. Remarkably, when feeding with visual inputs (e.g., white noise), simulated alpha oscillations propagated from lower level to higher level, while simulating resting-state gave rise to feedback waves.

The computational study suggests that the directionality of traveling waves could be closely linked with task conditions (visual processing vs. rest state), and is supported by human EEG studies where participants were instructed to monitor a visual luminance sequence or keep their eyes closed. However, those human experiments were conducted separately within different experimental sessions and different groups of participants, and it is thus difficult to infer a direct relationship between the task condition and waves’ direction. To verify the predictions of the computational work and to systematically examine how the waves’ direction changes from one task condition to another, the current EEG study was designed to incorporate stimulus-on periods (visual processing) and stimulus-off periods (resting state) within each single trial, by which we could trace the moment-to-moment changes of the waves’ direction caused by task conditions in a consistent way.

## Materials and Methods

### Participants

14 subjects participated in this experiment. One subject was rejected due to a technical problem during the experimental recording, leaving 13 subjects (6 females; mean age 25.57, range 21-31; 2 left-handed) for inclusion in the analysis. All participants reported no history of epileptic seizures or photosensitivity and they had normal or corrected to normal vision. Before starting the experiment, all participants gave written informed consent as specified by the Declaration of Helsinki. The study was performed under the guidelines for research according to author’s research institute at the “Centre de Recherche Cerveau et Cognition” and the protocol was approved by the committee “Comité de protection des Personnes Sud Méditerranée 1” (ethics approval number N° 2016-A01937-44).

### Stimuli generation

Visual stimuli were generated using MATLAB scripts and presented using the Psychophysics Toolbox(Brainard, 1997). The stimuli were displayed on a cathode ray monitor in a dark room, positioned 57 cm from the subjects, with a refresh rate of 160Hz and a resolution of 800 × 600 pixels. We used two types of visual luminance sequences (Figure 1) as visual inputs: dynamic (or white-noise) and static stimulation. For the white-noise sequences, the power spectrum was normalized to have equal power at all frequencies (up to 80 Hz). The resulting luminance of white-noise sequences ranged from black (0.1 cd/m^2^) to white (59 cd/m^2^), whereas the static ones were held constant with full contrast (59 cd/m^2^). Luminance sequences were displayed for 5 seconds within a disc of 3.5° radius which was centered at 7.5° above a center white dot on a black background.

**Figure 1.**
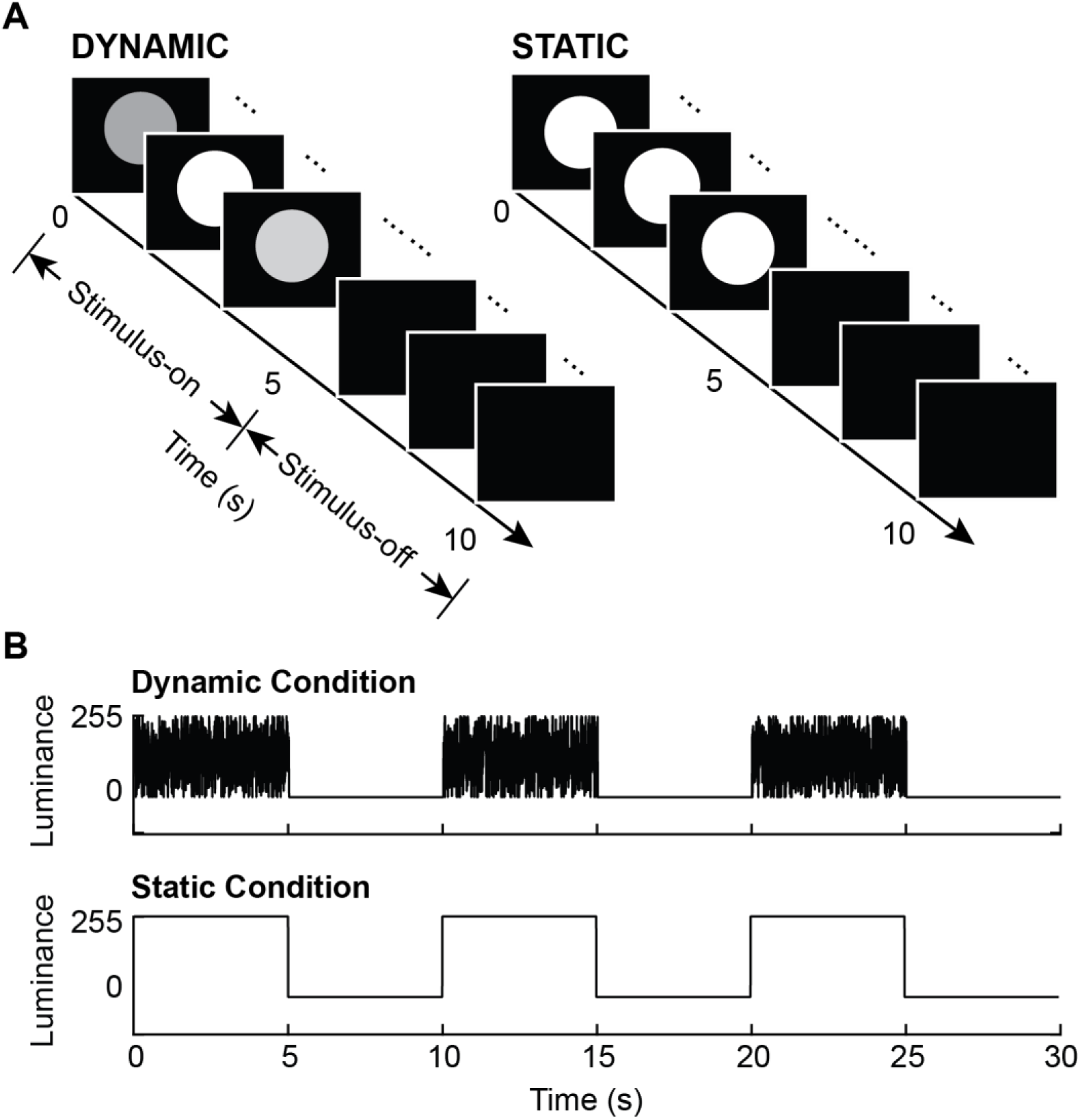
Experiment design. ***A.*** Two types of trials were included in this experiment. For static trials, the luminance of visual input was held constant at a value of 255 (full contrast), while for dynamic ones, the luminance changed randomly from 0 to 255 on each screen refresh. In both cases, luminance sequences were displayed for the first 5 seconds (Stimulus-on period), then followed by 5 seconds of blank screen (Stimulus-off period). ***B.*** Schematic diagrams of two subblocks for both dynamic and static conditions. Each subblock contained three identical trials, which made up a 30 s long time course.

### Experimental design

Subjects were instructed to perform a visual detection task. During the experiment, three identical trials (either static or dynamic) were displayed in a row, grouped into a subblock (Figure 1B). Prior to each subblock, a green center dot was displayed until subjects pressed the space bar to indicate their readiness. The green dot then disappeared and was followed by those three trials after a time-interval of 200∼300 ms. A prototypical trial started (Figure 1A) with 5 seconds of luminance sequences (either dynamic or static) in a disc above a white fixation dot at the center of the screen and then 5 seconds of blank screen. That is, each trial contained a stimulus-on period and a stimulus-off period, which allowed us to investigate the moment-to-moment changes of traveling waves when shifting from one task condition to another. Observers were asked to keep their fixation throughout the trial. Also, during visual stimulation (stimulus-on period), observers needed to covertly attend the disc to detect a brief square target (decreased luminance) inside the disc.

Two types of trials lead to two corresponding subblocks, dynamic or static which were presented alternatively and also counterbalanced within subjects. Targets (1 s) appeared at a random time (uniform distribution) from 0.25 s after the onset to 0.25 s before the offset of luminance sequences on a random 20% of trials. The square target luminance was adjusted according to each subject by a staircase procedure using the Quest function (Watson & Pelli, 1983) to ensure 80% detection rate. In dynamic trials, the luminance of the square target fluctuated according to the white-noise sequences, but with a lower contrast compared to the rest of the disk. The experiment was composed of five sessions of 10 experimental blocks of 6 trials (i.e., 2 subblocks) each, with a total duration of about 1 hour.

### EEG recording and Pre-processing

Continuous brain activity was recorded from the subjects using a 64-channel active BioSemi electroencephalography (EEG) system, with 1,024 Hz digitizing sample rate and 3 additional ocular electrodes. Custom scripts in the EEGlab toolbox (Delorme & Makeig, 2004) were applied to the pre-processing steps, during which both target-present and target-absent trials were included. We first rejected the noisy channels and then the data was offline down-sampled to 160 Hz. In order to remove power line artefacts, a notch filter (47-53 Hz) was applied. We applied an average-referencing and removed slow drifts by applying a high-pass filter (> 1 Hz). Data epochs were created around −0.5 s to 10 s around the trial onset, and EEG activity was corrected by subtracting the baseline activity from −0.25 s to 0 before trial onset. Finally, the data was screened manually for eye movements, blinks and muscular artefacts and whole epochs were rejected as needed.

### Wave quantification

In order to quantify the presence of traveling waves in EEG signals and assess the propagation direction, we adopted a wave quantification method from our previous studies (Alamia & VanRullen, 2019; Lozano-Soldevilla & VanRullen, 2019), which is described in Figure 3. For each subject, every trial (10 s long with 0.5 s baseline) was divided into 20 time-bins by a sliding window of 1 second (with 500 ms overlap). For each time-bin, we stacked EEG signals from seven midline electrodes (from posterior to frontal: Oz, POz, Pz, CPz, Cz, FCz, Fz) to form a 2D (electrode-time) map. To computationally quantify the waves’ amount, we utilized a 2D-FFT transform for each 2D map. This transform results in temporal frequencies along the horizontal axis as well as spatial frequencies along the vertical axis. The horizontal midline indicates stationary oscillations with no spatial propagation, while the upper and bottom quadrants reflect forward- and backward-propagating waves, respectively. We extracted the max value within the alpha-band temporal frequencies (8-13Hz) from the upper quadrant of the 2D-FFT as the FW value for this time window, and the max value (also within the alpha-band) from the lower quadrant as the BW value. After repeating this procedure over all 20 time-bins, we finally obtained two curves representing the dynamic changes of FW and BW waves along time.

To assess statistical significance of traveling waves, we used a non-parametric test. Specifically, we shuffled the electrodes’ order 100 times for each time bin, thereby eliminating any spatial organization of the oscillatory signals (including traveling waves). For this surrogate data set, we repeated the same 2D-FFT procedure as described above. Since the shuffling procedure only eliminated the spatial structure but left intact the oscillatory power of EEG signals, the resulting FW and BW curves (Figure 2C), based on the maximum power in each quadrant, could still fluctuate across time: oscillatory power was relatively suppressed during stimulus-on periods, then increased in the absence of visual input. These power fluctuations in the surrogate data, however, were similar in the FW and BW directions (as expected because of the shuffling procedure). In order to focus on the differences between real and surrogate data, we corrected the real wave patterns by dividing their values by the corresponding surrogate patterns, and expressing the result in dB units (Figure 2D).

**Figure 2.**
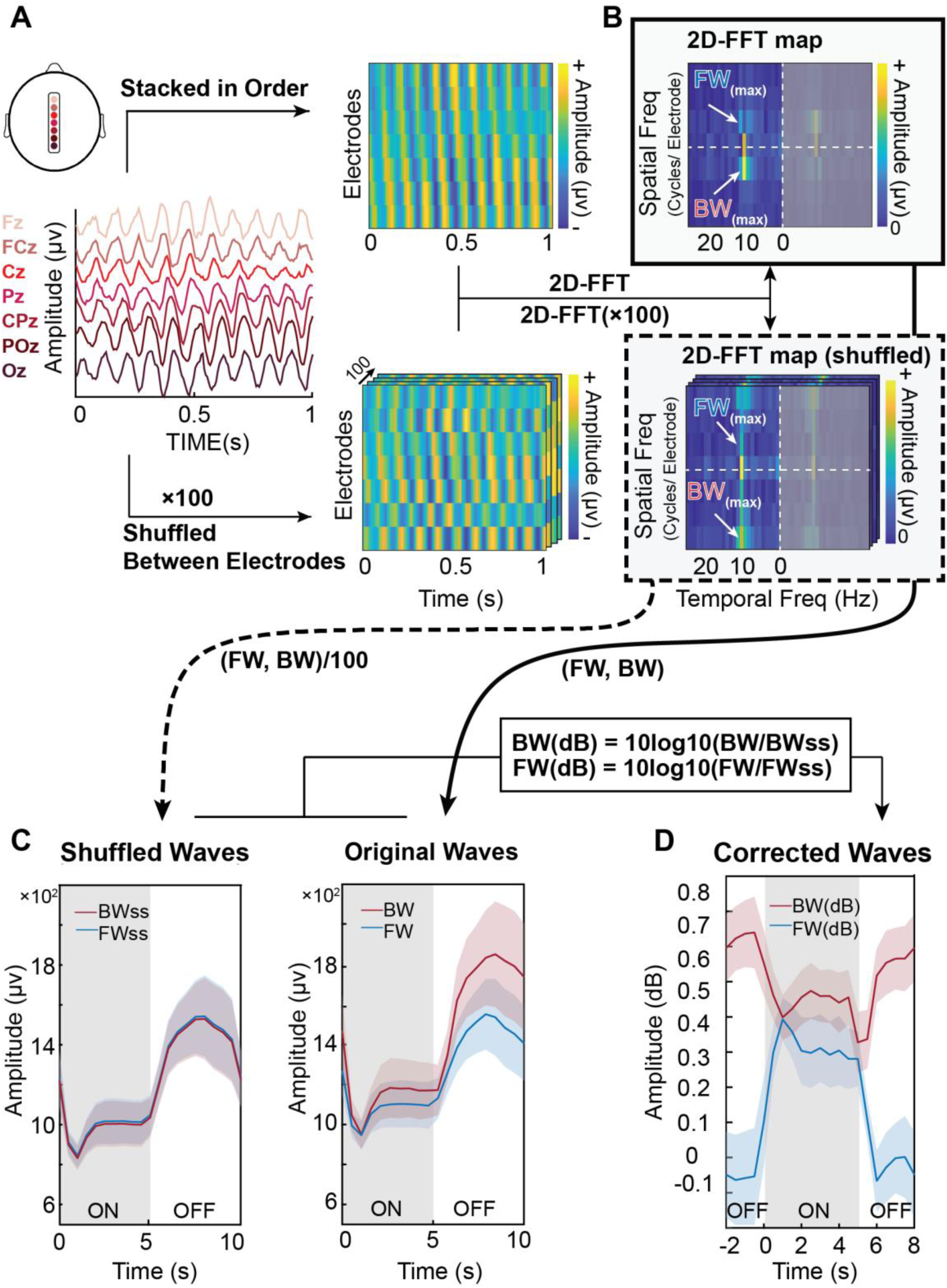
Alpha-band traveling waves in raw EEG signals. ***A. Left***: 7 midline electrodes of the 10-20 system are ordered from posterior to anterior (Oz to Fz) and backward-traveling waves (BW) can be observed in the 1 s long time window. ***Right***: A 2D-map of the same data with electrodes stacked in order and with amplitude color-coded (Top). To statistically quantify the waves’ direction, we employed a non-parametric test by shuffling the electrodes’ order for each time window 100 times. The resultant surrogate 2D-maps eliminate the spatial structure of the original signals, including their original propagating direction (Bottom). ***B.*** Temporal frequencies (x axis) and spatial frequencies (y axis) for both real and surrogate data are obtained by computing a 2D-FFT. The temporal frequencies were computed up to 80Hz, but only displayed until 25Hz for illustration purposes. Since the 2D-FFT gives symmetrical results around the origin, we only focused on the left part of the plot. The maximum value in the upper quadrant represents the strength of forward traveling waves, while the maximum value in the lower quadrant quantifies the strength of feedback traveling waves. ***C. Left***: For surrogate data, we averaged the 100 surrogate values separately for BW and FW signals and for each time bin. Shaded area stands for s.e.m across subjects. ***Right***: Similar time courses were obtained for the real data. ***D.*** The surrogate line plots were used as a baseline, mostly reflecting the background (alpha) oscillatory power. After correcting for these baseline fluctuations (and expressing the result in dB, as per the equations), we obtained a measure of the dynamics of FW and BW waves. Shaded area stands for s.e.m. across subjects; the stimulus-on period was shifted to the center part for better visualization of these dynamics around stimulus onset and offset.

**Figure 3.**
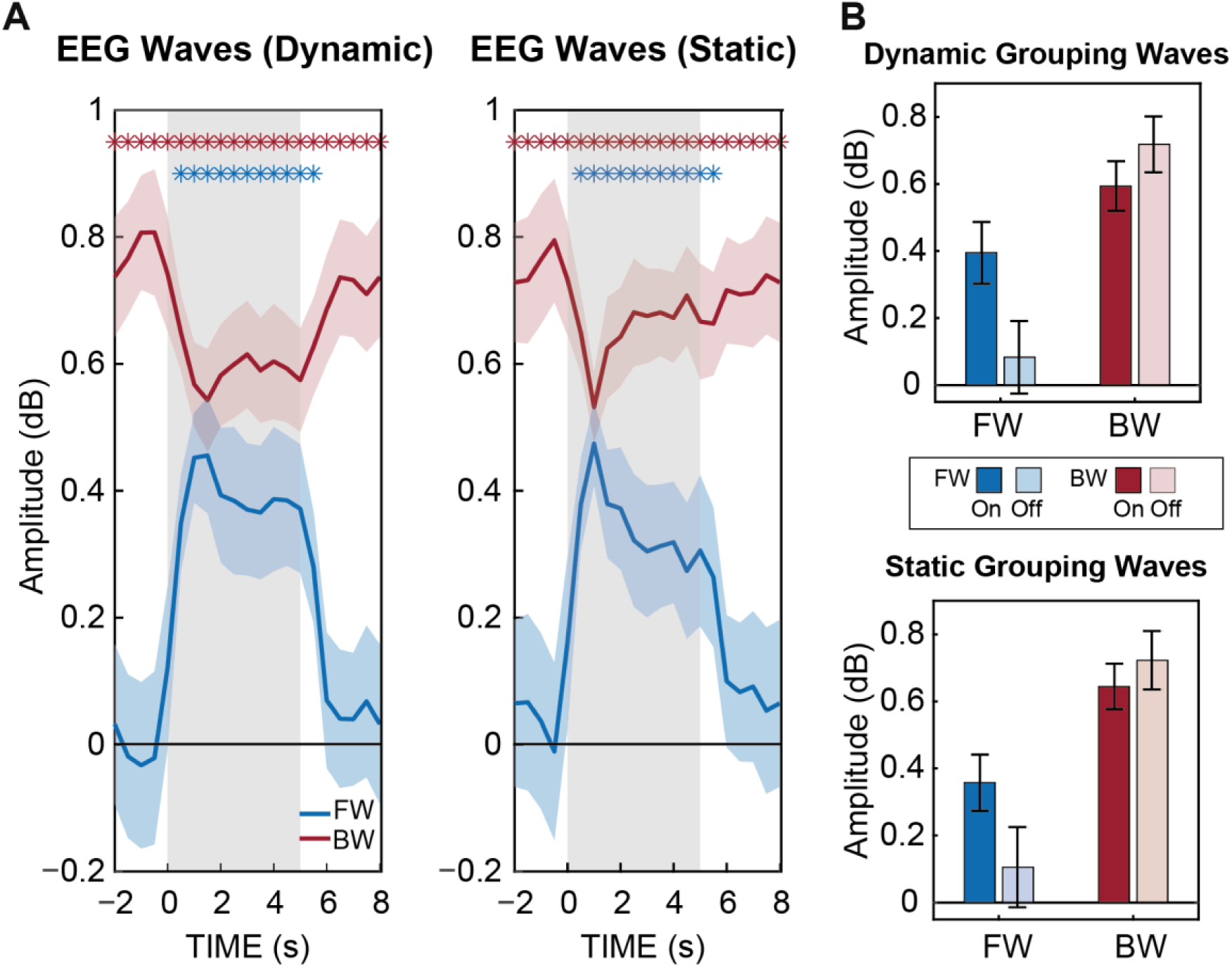
Dynamics of FW and BW waves over time under different task conditions and the corresponding bar graphs. ***A.*** The two plots show how waves evolve over time under dynamic (white-noise visual sequences) and static (full contrast visual sequences) task conditions. Blue and red asterisks represent separately the significant time points for FW waves (blue) and BW waves (red) when compared with zero. ***B.*** To better compare wave’ patterns, waves were grouped over time points within each period type (stimulus ON/OFF).

### Analysis

We first conducted 1-sample *t* tests against zero for both BW and FW waves separately to confirm their presence at each time point (corrected for multiple comparisons via FDR, α = 0.05). Second, we examined differences between the waves across the different experimental conditions. For this, we carried out a within-subject three-factor repeated measure ANOVA: CONDITIONS (static vs. dynamic visual stimulation) × WAVES (FW vs. BW) × TIME-BINS (20). To clearly examine the influence of tasks (visual processing vs. rest state), we also grouped all time points within the stimulus-on and stimulus-off periods and conducted another ANOVA with factors CONDITIONS (static vs. dynamic) × WAVES (FW vs. BW) × TASKS (stimulus-on vs. stimulus-off).

## Results

### Stimulus-evoked FW waves and spontaneous ongoing BW waves

Figure 3A illustrates the evolution of FW (blue) and BW (red) waves as a function of time under dynamic and static visual stimulation conditions, averaged across all subjects. The waves’ traces in both plots show similar patterns overall: BW waves are relatively high during both stimulus-on and stimulus-off periods (with a decrease during stimulation), while FW waves seem to only emerge after the onset of visual stimulation, and disappear after the offset of visual inputs. In other words, the occurrence of FW waves is highly dependent on external stimulation while BW waves exist both in the presence and absence of stimulation. Therefore, we propose that FW waves are associated with visual processing (e.g. as a stimulus-evoked wave), while BW waves reflect ongoing spontaneous or endogenous activity. To support this, we conducted 1-sample *t* tests against zero (p < 0.05, corrected for multiple comparisons by FDR) for the two waves separately at each time point. BW waves were significant during the entire time-course (significant values are marked with asterisks in Figure 3A). However, FW waves were only significant from 0.5 s to 5.5 s in both dynamic and static conditions.

### Both FW and BW waves are task-dependent

After establishing the presence of FW waves during visual stimulation, and BW waves during both visual stimulation and resting state, we further examined the properties of both waves under various task conditions, using a 3-way repeated measures ANOVA with factors CONDITIONS (dynamic/static), WAVES (FW/BW) and TIME-BINS (20 values). This revealed main effects for TIME-BINS (F_(19,228)_ = 19.083, p < 0.001) and WAVES (F_(1,12)_ = 7.048, p = 0.021), a significant two-way interactions for WAVES × TIME-BINS (F_(19,228)_ = 9.002, p < 0.001), as well as a significant three-way interaction (F_(19,228)_ = 3.103, p < 0.001).

The CONDITIONS × TIME-BINS interaction reached significance for FW waves (F_(19,228)_ = 2.698, p < 0.001) at time points 2 s, 3.5 s, and 5 s, and for BW waves (F_(19,228)_ = 2.928, p < 0.001) at time points 2∼4 s and 5 s. That is, the time-course of FW waves showed less power for static stimulation at certain time points. BW waves were also influenced by the stimulation type but with increasing power for static stimulation towards the later part of each stimulation period (Figure 3A). We speculate that both waves may be influenced by stimulus complexity since static stimuli are much simpler and more predictable compared to dynamic white-noise luminance sequences.

FW waves were only present during visual processing, while BW waves existed during both task conditions (visual processing vs. rest). To further examine whether BW waves showed significant differences associated with the tasks, another three-way repeated measures ANOVA was conducted with factors CONDITIONS (dynamic/static), WAVES (FW/BW) and TASKS (stimulus-on/off). That is, the TIME-BINS factor (20 values) was replaced with the TASKS factor (2 values). The wave amplitudes were averaged over time bins (separately for the stimulus-on and stimulus-off conditions). This time, we did not obtain a significant three-way interaction (Figure 3B). Instead, the ANOVA revealed a significant two-way interaction for WAVES × TASKS (F_(1,12)_ = 18.056, p = 0.001). As expected, the main effect of CONDITIONS was significant (F_(1,12)_ = 25.95, p < 0.001) for FW waves, similar to the result of 1-sample *t* tests above (Figure 3A). For BW waves, the main effect of CONDITIONS was also significant (F_(1,12)_ = 7.196, p = 0.02) with higher BW power in the absence of visual inputs. The modulation of BW waves by visual stimulation is in line with other studies showing that spontaneous traveling waves could be suppressed by external inputs (Patten et al., 2012; Sato et al., 2012).

In summary, FW waves appear to be caused by external visual stimulation, while BW waves can originate spontaneously but could be reduced (yet not eliminated) by the presence of visual stimulation. During visual stimulation, both waves are present and modulated by the type of visual inputs, with lower FW but higher BW waves’ power for simpler (static) sensory stimulation.

### FW and BW waves are negatively related during visual stimulation

During visual stimulation (stimulus-on periods), both FW and BW waves appear to be simultaneously present (Figure 3A). However, the average traveling waves behavior does not necessarily reflect the instantaneous state of the brain and its dynamics: FW and BW waves may by truly equally present at each moment in time, or they may tend to happen in alternation, at distinct moments in time. To further examine the relationship between them, we assessed the moment-to-moment correlation between FW and BW waves (Figure 4A) for each stimulus condition. Figure 4A shows scatter plots from a representative subject. For each condition, each 1 s time-bin window produces one pair of FW and BW wave values. To discard the common influence of oscillatory amplitude fluctuations on both FW and BW traveling waves, we correct each wave’s value by its corresponding averaged surrogate value (obtained by shuffling the electrodes’ order 100 times, as explained in Figure 2A). The correlation between the resulting FW and BW values in dB units showed a clear and significant (p < 0.01) negative trend, for both the dynamic and static stimulus conditions. This means that when FW waves were stronger, BW waves tended to be weaker, and vice-versa. Figure 4B gives the average correlation across all subjects: significant negative correlation between FW and BW waves can be observed for both the dynamic (mean = −0.732 ± 0.053, t = −49.371, p < 0.001) and static conditions (mean = −0.74 ± 0.056, t = −47.644, p < 0.001).

**Figure 4.**
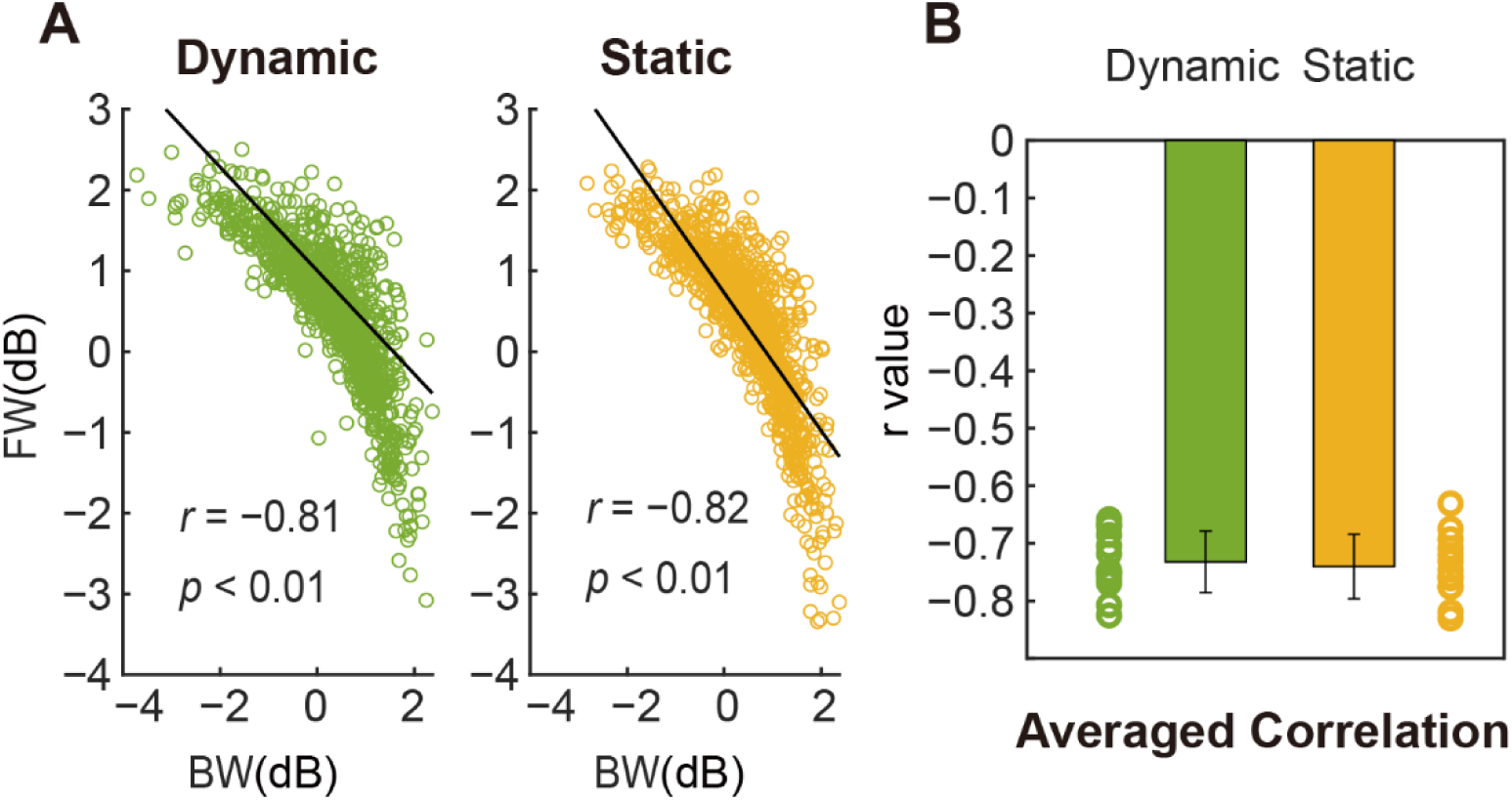
FW and BW waves are negatively correlated during stimulus-on periods. **A.** Results from a representative subject. The two scatter plots show the correlation between corrected BW and FW values under dynamic stimulation (left) and static stimulation (right). Black lines are regression lines. **B.** Bar graphs show averaged correlation (r values) across all subjects for dynamic (green) and static (yellow) conditions. The corresponding circles are individual results.

## Discussion

Based on EEG data from human participants, we demonstrated that the direction of alpha traveling waves (8-13Hz) is task-dependent, confirming suggestions from prior studies (Alamia & VanRullen, 2019; Halgren et al., 2019; Lozano-Soldevilla & VanRullen, 2019; Zhang et al., 2018), and verifying the predictions of our own modeling study on the generation and propagation of alpha oscillations (Alamia & VanRullen, 2019). Specifically, we characterized FW waves traveling from occipital to parietal regions elicited by visual stimulation, and BW waves in the reversed direction dominating during rest state. Furthermore, the presence of external visual stimulation reduced BW waves (Figure 3), which is in line with other studies on spontaneous traveling waves (Patten et al., 2012; Sato et al., 2012). Lastly, during visual stimulation, FW waves and BW waves coexisted on average, but were in fact negatively correlated over time.

### Contributions of the current study

It should be emphasized that the current experimental design directly contrasted the conditions of visual processing and resting state within each trial. Previously, a number of studies had examined traveling waves under various single-task conditions, including visual stimulation (Alamia & VanRullen, 2019; Lozano-Soldevilla & VanRullen, 2019; Muller et al., 2014; Nauhaus et al., 2012), sleep (Muller et al., 2016) or quiet wakefulness (Alamia & VanRullen, 2019; Halgren et al., 2019). While these experiments confirmed the existence of traveling waves, they did not make it possible to track how the waves change from one condition to another. Due to the within-subject design in the present study, we found that the waves’ direction is highly sensitive to the task conditions.

Compared with previous studies using dynamic white noise sequences as visual stimulation (Alamia & VanRullen, 2019; Lozano-Soldevilla & VanRullen, 2019), we also included a simpler type of visual stimulation: static luminance sequences. The results showed that although these two stimulus types evoked similar FW and BW waves, towards the later stages of visual stimulation, the BW wave power increased at time points 2∼4 s, 5s and FW wave power decreased at 2 s, 3.5 s, and 5 s. This may be due to the relative simplicity of static inputs compared to the dynamic ones.

Unlike prior studies measuring a single traveling wave direction from phase gradients over certain brain regions (Halgren et al., 2019; Zhang et al., 2018), we here derived two opposite components of the waves’ direction from the pattern of brain activity within each time window, and quantified their strength. Previous studies have shown that traveling waves can propagate in different directions (Alexander et al., 2009; Massimini et al., 2004), and that the co-existence of two opposite waves may cause a loss of wave information (Alexander et al., 2013). For example, under synchronized states like sleep or resting-state, traveling waves have often been reported to propagate in a frontal-to-occipital direction (Massimini et al., 2004). However, traveling waves are less frequently observed under more complex cognitive states (Alexander et al., 2013; Muller et al., 2014), this may be caused by the interference of waves propagating in opposite directions, while their direction is characterized as a single value. Instead, our analysis method independently quantifies waves propagating in the two directions. Also, the separation of FW and BW waves’ components contributed to reveal their distinct functional roles. We revealed a closer link between FW waves and visual processing as an evoked wave, since FW waves emerged at the onset of visual input and disappeared right after the offset (Figure 3); meanwhile, BW waves were more related to the resting state, acting as a spontaneous wave.

### An explanation under the predictive coding framework

The generation and directionality of traveling waves can tentatively be interpreted within the predictive coding framework (Rao & Ballard, 1999). In our previous work (Alamia & VanRullen, 2019), we built a seven-level hierarchical model of visual cortex with bidirectional connectivity implementing predictive coding. Within the hierarchy, higher levels predicted the activity of lower ones through inhibitory feedback, and lower levels sent the prediction error via feedforward excitation to the higher layers in order to correct their prediction. With biologically plausible parameters (neural time constants, communication delays), this model produced alpha rhythms traveling through the hierarchy. The waves could travel in the FW direction when the model was presented with visual inputs, and in the BW direction in the absence of inputs (while the model was processing “top-down priors” instead of bottom-up sensory signals).

In this context, it is reasonable to infer that FW waves carry “residual error” signals (the difference between the actual visual inputs and the prediction from higher-level regions), while BW waves carry the prediction signals. Remarkably, the current results that FW waves emerged only during visual stimulation and BW waves were dominant in the resting state agree with this framework. On the other hand, the negative correlation across time between FW waves and BW waves during visual stimulation may reflect the dynamics of predictive coding mechanism. That is, stronger prediction signals within BW waves are associated with weaker prediction errors carried by FW waves and vice versa. Moreover, in the static condition, BW waves increased but FW waves decreased significantly at the later stages of visual stimulation, indicating that prediction information becomes stronger while error signals weaken over time. This was not the case in the dynamic condition, which has much more complex (and unpredictable) stimulus temporal structure, leading to less precise prediction signals and larger error signals.

### Spontaneous BW waves may reveal top-down control

Spontaneous ongoing waves have been reported in the cortex under anesthesia or quiet wakefulness (Alamia & VanRullen, 2019; Petersen et al., 2003; Sakata & Harris, 2009). The current study points to BW waves as spontaneous waves, given their existence under resting state. Besides, the significant reduction of BW waves due to the presence of visual inputs also agrees with other studies on spontaneous traveling waves (Patten et al., 2012; Sato et al., 2012). This reduction could be explained by the desynchronization caused by visual processing, since spontaneous activity measured during quiet wakefulness may reflect synchronized cortical states (Harris & Thiele, 2011). On the other hand, given the spatial extent of traveling waves across distributed cortical regions, their functional role may entail long-range information integration (Halgren et al., 2019; Sato et al., 2012). In particular, it is speculated that BW waves may participate in the organization of top-down or feedback information flow (Halgren et al., 2019; Van Kerkoerle et al., 2014). This is in line with the dominance of alpha-band activity in the waves, a frequency which is typically associated with top-down control (Bahramisharif et al., 2013; Foxe & Snyder, 2011; Jensen et al., 2012; Klimesch, Sauseng, et al., 2007). A concrete example of the top-down control by traveling waves comes from a recent study. Davis and colleagues (Davis et al., 2019) reported that spontaneous traveling cortical waves modulate perception by a gating mechanism: as waves propagate across the cortex, the brain excitability is periodically altered into successive depolarizing or hyperpolarizing states which then cause an increase or decrease (respectively) in perceptual sensitivity to incoming inputs.

### Stimulus-evoked FW waves are associated with bottom-up sensory processing

In the current study, we measured alpha FW waves which were directly linked with visual processing (and absent during rest). This direct link is also supported by our prior studies of perceptual echoes: since these echoes are measured by cross-correlation with the visual input sequence, they can be viewed as a direct reflection of visual processing (Alamia & VanRullen, 2019; Lozano-Soldevilla & VanRullen, 2019; VanRullen & Macdonald, 2012). Recent work from our group found that these perceptual echoes propagate from occipital to parietal regions in a forward direction (Alamia & VanRullen, 2019; Lozano-Soldevilla & VanRullen, 2019). Although further research is needed to test whether FW waves also contribute to sensory processing in other modalities (like audition or touch), FW waves may serve to integrate the information flow along the bottom-up path. This is consistent with the “scanning hypothesis” proposed by (Pitts & McCulloch, 1947), suggesting that the alpha rhythm repeatedly scans the visual cortex. The bidirectionality of alpha traveling waves found in the current study may help to clarify an apparent contradiction between the conventionally postulated inhibitory role of alpha oscillations (Bonnefond & Jensen, 2012; Jensen & Mazaheri, 2010), and their reported implication in sensory processing (VanRullen, 2016; Varela et al., 1981). Inhibition may be carried by the BW component of alpha oscillations as mentioned above, whereas, the FW component may reflect the positive relation between alpha and sensory processing.

### Conclusion

In summary, the current study corroborated the predictions from our prior EEG and modelling study (Alamia & VanRullen, 2019). It showed that FW and BW waves are inversely related to sensory processing, and may characterize opposite directions of information flow in the brain hierarchical system. Importantly, the transitions between FW and BW waves were observed within single trials and for the same human subjects. First, FW waves travel from occipital to frontal regions during visual processing, while BW waves are spontaneously generated and travel in the opposite direction, likely reflecting a feedback process. Second, during visual stimulation, both FW and BW waves exist on average, but are negatively correlated across time, suggesting that they reflect distinct functions that may draw on common brain resources.

## Notes

### Competing Interest Statement

The authors have declared no competing interest.

### Summary of Updates

Section on "Both FW and BW waves are task-dependent" updated to show the influence of stimulation type on FW waves; Figure 4 revised.

## References

Alamia, A., & VanRullen, R. (2019). Alpha oscillations and traveling waves : Signatures of predictive coding ? PLOS Biology, 1–26. https://doi.org/10.17605/OSF.IO/NC4RG

Alexander, D. M., Flynn, G. J., Wong, W., Whitford, T. J., Harris, A. W. F., Galletly, C. A., & Silverstein, S. M. (2009). Spatio-temporal EEG waves in first episode schizophrenia. Clinical Neurophysiology, 120(9), 1667–1682. https://doi.org/10.1016/j.clinph.2009.06.020

Alexander, D. M., Jurica, P., Trengove, C., Nikolaev, A. R., Gepshtein, S., Zvyagintsev, M., Mathiak, K., Schulze-Bonhage, A., Ruescher, J., Ball, T., & van Leeuwen, C. (2013). Traveling waves and trial averaging: The nature of single-trial and averaged brain responses in large-scale cortical signals. NeuroImage, 73, 95–112. https://doi.org/10.1016/j.neuroimage.2013.01.016

Bahramisharif, A., van Gerven, M. A. J., Aarnoutse, E. J., Mercier, M. R., Schwartz, T. H., Foxe, J. J., Ramsey, N. F., & Jensen, O. (2013). Propagating neocortical gamma bursts are coordinated by traveling alpha waves. Journal of Neuroscience, 33(48), 18849–18854. https://doi.org/10.1523/JNEUROSCI.2455-13.2013

Bonnefond, M., & Jensen, O. (2012). Report Alpha Oscillations Serve to Protect Working Memory Maintenance against Anticipated Distracters. Current Biology, 22(20), 1969–1974. https://doi.org/10.1016/j.cub.2012.08.029

Brainard, D. H. (1997). The Psychophysics Toolbox. Spatial Vision, 10(4), 433–436.

Davis, Z. W., Muller, L., Trujillo, J., Sejnowski, T., & Reynolds, J. H. Spontaneous Traveling Cortical Waves Gate Perception in Awake Behaving Primates. bioRxiv 811471 [Preprint]. 2019 [cited 2020 Apr 15]. Available from: https://www.biorxiv.org/content/10.1101/811471v1

Delorme, A., & Makeig, S. (2004). EEGLAB : an open source toolbox for analysis of single-trial EEG dynamics including independent component analysis. 134, 9–21. https://doi.org/10.1016/j.jneumeth.2003.10.009

Ermentrout, G. B., & Kleinfeld, D. (2001). Traveling Electrical Waves in Cortex. Neuron, 29(1), 33–44. https://doi.org/10.1016/s0896-6273(01)00178-7

Fellinger, R., Gruber, W., Zauner, A., Freunberger, R., & Klimesch, W. (2012). Evoked traveling alpha waves predict visual-semantic categorization-speed. NeuroImage, 59(4), 3379–3388. https://doi.org/10.1016/j.neuroimage.2011.11.010

Fiser, J., Chiu, C., & Weliky, M. (2004). Small modulation of ongoing cortical dynamics by sensory input during natural vision. Nature, 431(7008), 573–578. https://doi.org/10.1038/nature02907

Foxe, J. J., & Snyder, A. C. (2011). The role of alpha-band brain oscillations as a sensory suppression mechanism during selective attention. Frontiers in Psychology, 2(JUL), 1–13. https://doi.org/10.3389/fpsyg.2011.00154

Halgren, M., Ulbert, I., Bastuji, H., Fabó, D., Eross, L., Rey, M., Devinsky, O., Doyle, W. K., Mak-McCully, R., Halgren, E., Wittner, L., Chauvel, P., Heit, G., Eskandar, E., Mandell, A., & Cash, S. S. (2019). The generation and propagation of the human alpha rhythm. Proceedings of the National Academy of Sciences of the United States of America, 116(47), 23772–23782. https://doi.org/10.1073/pnas.1913092116

Harris, K. D., & Thiele, A. (2011). Cortical state and attention. Nature Reviews Neuroscience, 12(9), 509–523. https://doi.org/10.1038/nrn3084

Jensen, O., Bonnefond, M., & VanRullen, R. (2012). An oscillatory mechanism for prioritizing salient unattended stimuli. Trends in Cognitive Sciences, 16(4), 200–206. https://doi.org/10.1016/j.tics.2012.03.002

Jensen, O., & Mazaheri, A. (2010). Shaping functional architecture by oscillatory alpha activity: gating by inhibition. Front. Hum. Neurosci, 4, 1–8. https://doi.org/10.3389/fnhum.2010.00186

Klimesch, W., Hanslmayr, S., Sauseng, P., Gruber, W. R., & Doppelmayr, M. (2007). P1 and traveling alpha waves: Evidence for evoked oscillations. Journal of Neurophysiology, 97(2), 1311–1318. https://doi.org/10.1152/jn.00876.2006

Klimesch, W., Sauseng, P., & Hanslmayr, S. (2007). EEG alpha oscillations : The inhibition – timing hypothesis. Brain Research Reviews, 53, 63–88. https://doi.org/10.1016/j.brainresrev.2006.06.003

Lozano-Soldevilla, D., & VanRullen, R. (2019). The Hidden Spatial Dimension of Alpha : 10-Hz Perceptual Echoes Propagate as Periodic Traveling Waves in the Human. CellReports, 26(2), 374–380. https://doi.org/10.1016/j.celrep.2018.12.058

Massimini, M., Huber, R., Ferrarelli, F., Hill, S., & Tononi, G. (2004). The sleep slow oscillation as a traveling wave. Journal of Neuroscience, 24(31), 6862–6870. https://doi.org/10.1523/JNEUROSCI.1318-04.2004

Muller, L., Chavane, F., Reynolds, J., & Sejnowski, T. J. (2018). Cortical traveling waves: Mechanisms and computational principles. Nature Reviews Neuroscience, 19(5), 255–268. https://doi.org/10.1038/nrn.2018.20

Muller, L., Piantoni, G., Koller, D., Cash, S. S., Halgren, E., & Sejnowski, T. J. (2016). Rotating waves during human sleep spindles organize global patterns of activity that repeat precisely through the night. ELife, 5, 1–16. https://doi.org/10.7554/elife.17267

Muller, L., Reynaud, A., Chavane, F., & Destexhe, A. (2014). The stimulus-evoked population response in visual cortex of awake monkey is a propagating wave. Nature Communications, 5, 1–14. https://doi.org/10.1038/ncomms4675

Nauhaus, I., Busse, L., Ringach, D. L., & Carandini, M. (2012). Robustness of traveling waves in ongoing activity of visual cortex. Journal of Neuroscience, 32(9), 3088–3094. https://doi.org/10.1523/JNEUROSCI.5827-11.2012

Patten, T. M., Rennie, C. J., Robinson, P. A., & Gong, P. (2012). Human cortical traveling waves: Dynamical properties and correlations with responses. PLoS ONE, 7(6), e38392. https://doi.org/10.1371/journal.pone.0038392

Petersen, C. C. H., Hahn, T. T. G., Mehta, M., Grinvald, A., & Sakmann, B. (2003). Interaction of sensory responses with spontaneous depolarization in layer 2/3 barrel cortex. Proceedings of the National Academy of Sciences of the United States of America, 100(23), 13638–13643. https://doi.org/10.1073/pnas.2235811100

Pitts, W., & McCulloch, W. S. (1947). How we know universals the perception of auditory and visual forms. The Bulletin of Mathematical Biophysics, 9(3), 127–147. https://doi.org/10.1007/BF02478291

Rao, R. P. N., & Ballard, D. H. (1999). Predictive coding in the visual cortex: A functional interpretation of some extra-classical receptive-field effects. Nature Neuroscience, 2(1), 79–87. https://doi.org/10.1038/4580

Sakata, S., & Harris, K. D. (2009). Population Activity in Auditory Cortex. Neuron, 64(3), 404–418. https://doi.org/10.1016/j.neuron.2009.09.020.

Sato, T. K., Nauhaus, I., & Carandini, M. (2012). Traveling Waves in Visual Cortex. Neuron, 75(2), 218–229. https://doi.org/10.1016/j.neuron.2012.06.029

Van Kerkoerle, T., Self, M. W., Dagnino, B., Gariel-Mathis, M. A., Poort, J., Van Der Togt, C., & Roelfsema, P. R. (2014). Alpha and gamma oscillations characterize feedback and feedforward processing in monkey visual cortex. Proceedings of the National Academy of Sciences of the United States of America, 111(40), 14332–14341. https://doi.org/10.1073/pnas.1402773111

VanRullen, R. (2016). Perceptual Cycles. Trends in Cognitive Sciences, 20(10), 723–735. https://doi.org/10.1016/j.tics.2016.07.006

VanRullen, R., & Macdonald, J. S. P. (2012). Perceptual Echoes at 10 Hz in the Human Brain. Current Biology, 22, 995–999. https://doi.org/10.1016/j.cub.2012.03.050

Varela, F. J., Toro, A., Roy John, E., & Schwartz, E. L. (1981). Perceptual framing and cortical alpha rhythm. Neuropsychologia, 19(5), 675–686. https://doi.org/10.1016/0028-3932(81)90005-1

Watson, A. B., & Pelli, D. G. (1983). QUEST: A general multidimensional bayesian adaptive psychometric method. Journal of Vision, 33(2), 113–120. https://doi.org/10.1167/17.3.10

Zhang, H., Watrous, A. J., Patel, A., Jacobs, J., Zhang, H., Watrous, A. J., Patel, A., & Jacobs, J. (2018). Theta and Alpha Oscillations Are Traveling Waves in the Human Neocortex Article Theta and Alpha Oscillations Are Traveling Waves in the Human Neocortex. Neuron, 98(6), 1269–1281. https://doi.org/10.1016/j.neuron.2018.05.019

